# Profilin1 regulates triple negative breast cancer cells’ migration through stabilization of Angiomotin and thereby YAP nuclear translocation

**DOI:** 10.64898/2026.08.28.747794

**Authors:** Chandana Praneetha Vipparthy, Sunil Kumar Manna

## Abstract

The Hippo pathway effector YAP1 is a potent oncogenic driver in triple-negative breast cancer (TNBC) and its activity is restrained by the scaffold protein Angiomotin-p130 (AMOT). AMOT is itself short-lived, being targeted for proteasomal degradation by NEDD4-family E3 ubiquitin ligases that dock at its L/P-PxY motifs. Here we identify Profilin1 (PFN1), an actin-binding protein with established actin-independent tumour-suppressive signalling functions in TNBC as a direct binding partner and stabilizer of AMOT. PFN1 and AMOT are co-immunoprecipitated, they share 70 common interactors and NEDD4 is one of them. Protein-protein docking shows the interaction of PFN1 on the first PPxY motif of AMOT, through its actin-binding domain. We further show that PFN1’s binding leaves the AMOT LPTY motif and both coiled-coil domains entirely unoccupied. Site-directed mutagenesis of AMOT PPxY motifs shows that PFN1 binding is unaffected by substitution of the PPxY tyrosines Y242 and Y287, either alone or in combination, indicating that PFN1 engages through its actin-binding domain. Functionally, PFN1 stabilizes AMOT as shown by cycloheximide-chase assay in TNBC. PFN1 induction increases cytoplasmic retention of YAP1, reduces TEAD occupancy at the CTGF promoter and thereby suppresses TNBC cell migration. Thus, this study suggests that PFN1 deregulates tumour cells migration by interacting with AMOT through its actin-binding domain, stabilizing AMOT and thereby arresting YAP in the cytoplasm, which might be an important therapeutic target to regulate TNBC.

## Introduction

Malignant tumours are characterized by their ability to invade and metastasize to distant normal tissues. In this intricate and highly selective process, tumour cells break away from their primary site and disseminate by various routes, such as the blood and lymph vessels, and reach hospitable ectopic site and grow in this new niche. During these steps, the ability of cells to migrate and proliferate are essential. Cell migration is a core component of tumour cell invasion and metastasis, which involves dynamic re-modelling of actin cytoskeleton through the coordinated actions of many different classes of actin-binding proteins (ABPs) such as Gelsolin, Profilin, VASP, N-WASP, Cofilin, etc. (Pollard and Borisy, 2003; Small et al., 2002). Altered expression of these ABPs have been linked to changes in actin cytoskeleton observed in malignant cells (Yamaguchi and Condeelis, 2007).

Breast tumour is one of the major cancers in women that causes morbidity world-wide (Bray et al., 2024), especially, triple-negative breast cancer (TNBC) cells that lack expression of the estrogen receptor, progesterone receptor and HER2, and consequently cannot be treated with the endocrine- or HER2-directed therapies available for other breast cancer subtypes (Bergin and Loi, 2019). Targeted options remain limited, and conventional chemotherapies are still the main treatment and outcomes frequently lead to relapse and metastasis (Bergin and Loi, 2019). Identifying the signalling molecules that drive TNBC invasion, and regulation of the endogenous molecules, which might give a brake in these events, are in need today to control the TNBC.

Profilin1 is a small (15 kDa) protein having binding domain with actin (ABD), phosphatidyl inositol (PtdIns-BD) and poly-L-proline (PLP-BD) (Ding et al., 2012). It is a ubiquitously expressed G-actin-binding protein in a stable 1:1 complex and acts as a nucleotide exchange factor, by catalyzing ADP-to-ATP exchange on G-actin and shuttling ATP-G-actin to the barbed ends of actin filaments (Witke, 2004). Apart from regulating the dynamics of actin filament turnover, Profilin functions as a hub that controls a complex network of molecular interactions involved in membrane trafficking (Dynamin1, Clathrin, etc.), focal contacts (VASP, Mena), cellular signalling (Rac/Rho pathway) and nuclear translocation (Exportin6, SMN) (Ding et al., 2012; Stüven et al., 2003). PFN1 expression is reduced in breast tumours, its silencing enhances the motility and invasiveness of MDA-MB-231 cells, and its restoration suppresses tumorigenicity and micrometastasis in xenografts (Zou et al., 2007). PFN1 overexpression attenuates AKT activation by stabilizing PTEN (Das et al., 2009), and regulates PI(3,4)P2 and lamellipodin at the leading edge (Bae et al., 2010). Work from our laboratory has extended this actin-independent repertoire in the same MDA-MB-231 model used here: PFN1 associates with PTEN and suppresses NF-κB activation by blocking IKK-dependent IκBα degradation (Zaidi and Manna, 2016); PFN1 induction, whether ectopic or driven by all-trans retinoic acid (ATRA), potentiates chemotherapeutic-agent-induced tumour regression (Saurav and Manna, 2022a); and PFN1 physically interacts with and stabilizes AMPKα, driving mTOR inhibition and autophagy (Saurav and Manna, 2022b). A recurring theme across these studies is that PFN1 acts by binding a partner protein and altering its stability or activity.

One of the cascades regulating organ size and tissue growth is the Hippo pathway, which acts by phosphorylating and inactivating the transcriptional co-activators YAP and TAZ (Yu et al., 2015). When Hippo signalling is low, unphosphorylated YAP accumulates in the nucleus and partners with TEAD transcription factors to drive proliferative and pro-migratory target genes such as CTGF (CCN2) and CYR61 (Zhao et al., 2011). In TNBC, YAP1 behaves as an oncogenic driver: it is preferentially nuclear, correlates with an epithelial-to-mesenchymal transition (EMT) signature, and its silencing impairs proliferation, migration and invasion (Andrade et al., 2017).

The scaffold protein Angiomotin helps control YAP alongside the core Hippo kinase cascade, and it does so through at least three distinct mechanisms. First, AMOT tethers YAP directly: its N-terminal region carries an LPTY motif and a PPxY motif (residues 239-242) that dock into the WW domains of YAP, sequestering it in the cytoplasm and repressing TEAD-dependent transcription (Chan et al., 2011; Zhao et al., 2011; Yi et al., 2013). Second, AMOT binds YAP through its coiled-coil (CC) domain, independently of the PY motifs (Moleirinho et al., 2014). Third, and mechanistically distinct from both, the AMOT CC domain scaffolds and activates LATS1/2, which phosphorylates YAP at S127 to create a 14-3-3 binding site and thereby drives cytoplasmic retention without requiring YAP to remain physically bound to AMOT at all (Paramasivam et al., 2011). The second PPxY (284-287), serves principally as a docking site for KIBRA rather than YAP (Paramasivam et al., 2011; Moleirinho et al., 2014). AMOT abundance is itself tightly controlled. NEDD4-family HECT-domain E3 ubiquitin ligases (NEDD4, NEDD4-2 and Itch) bind the same L/P-PxY motifs used for YAP engagement and target AMOT- p130 for poly-ubiquitination and proteasomal degradation, competing with YAP for access to the same surface (C. Wang et al., 2012). AMOT protein levels, and its capacity to restrain YAP, are therefore jointly governed at a single proline-motif-centred node - a node not previously linked to any actin-binding protein.

To further explore the role of PFN1, we performed mass spectrometry of PFN1, and found a number of new possible interacting candidates. Among them, we have chosen Angiomotin for further studies. AMOT is reported as a novel component that inhibits Yes-associated protein (YAP) oncoprotein of the Hippo pathway (Zhao et al., 2011). The pathway is composed of upstream regulators, central core kinase complexes, and downstream effectors: Yes Associated proteins (YAP) and transcriptional coactivator with PDZ-binding motif (TAZ), which further transcribe the TEAD genes (Yu et al., 2015). Recent studies have shown that hippo pathway also plays a prominent role in mediating angiogenesis in cancer cells (Boopathy and Hong, 2019). Hence, it needs to be studied if Profilin1 is in complex with AMOT and YAP, and if so, which domain of either AMOT or YAP is interacting with Profilin1. AMOT p130, AMOTL1 and AMOTL2 have conservative coiled-coil domains and one C-terminal PDZ domain (Moleirinho et al., 2014). Profilin1 is known to have a Poly-L-Proline-binding domain, which has high affinity towards proline-rich sequences, including PPXY-type motifs (Ding et al., 2009). This provides a hypothesis of Profilin competing with YAP to bind with AMOT and thereby play a role in regulating YAP/AMOT.

## Materials and Methods

### Cell culture

Cell lines used for this study: MDA-MB-231 and HEK293T. Cells were obtained from American Type Culture Collection (ATCC) (VA, USA). All the cells were grown in DMEM supplemented with 10% heat inactivated FBS, L-glutamine (2 mM), penicillin (100 U/ml), and streptomycin (100 µg/ml) and maintained in 5% CO_2_ at 37 °C. Cells were used from passage number 8 and experiments were carried out till passage number 15. Mycoplasma contamination-free cells were used as detected by the Gen-Probe Mycoplasma Rapid Detection kit (Fisher Scientific, Pittsburgh, PA, USA).

### Reagents and antibodies

NP40, NaCl, DMSO, EDTA, Tris, Glycine, all-trans retinoic acid (ATRA), Cycloheximide (CHX) and proteosomal inhibitor MG-132 were purchased from Sigma Aldrich Chemicals (St Louis, MO, USA). Sphingosine-1-phosphate (S1P) was purchased from MedChemExpress (Monmouth Junction, NJ, USA). Antibodies against AMOT, YAP and profilin 1 were obtained from Cell Signaling Technologies (Danvers, MA, USA). Antibodies against AMOT (raised in mice), HA, GAPDH and Steptavidin beads were obtained from Santa Cruz Biotechnology (Santa Cruz, CA, USA). Penicillin, streptomycin, DMEM, fetal bovine serum (FBS) (US origin) and Lipofectamine 2000 were obtained from Life Technologies (Grand Island, NY). Horseradish peroxidase (HRP)-conjugated - anti-rabbit IgG (Cat# 1140380011730) and - anti-mouse IgG (Cat# 1140680011730) antibodies were purchased from GeNei (Bengaluru, India).

### Plasmids and stable cell generation

HA-AMOT, HA-AMOT Y242A, HA-AMOT Y287A and HA-AMOT Y242A/Y287A were purchased from Addgene (Watertown, MA). SFB-PFN1 plasmid and MDA-MB-231 PFN1 stable cells were generated by Adeel H Zaidi as described previously (Zaidi and Manna, 2016). Angiomotin stable cells were established using MDA-MB-231 cells, designated as MDA-MB- 231 (AMOT^+/+^) (Amot-stable) in the laboratory as described previously (Zaidi and Manna, 2016).

### Immunoblotting

After treatment, whole cell extracts (WCE) were prepared using NETN lysis buffer (20 mM Tris-Cl pH 8.0, 2 mM EDTA, 250 mM NaCl and 0.1% NP40) with protease inhibitors. 50 and 100 µg of total protein from WCE were resolved on 12.5% SDS-PAGE. Resolved proteins were transferred onto PVDF membrane using wet transfer method at 90V for 3 h. After 3 h of transfer in cold condition, membrane was washed with TBST and incubated in 5% blocking buffer [5% (w/v) skimmed milk powder in TBST] to suppress non-specific IgG binding. After blocking, immunoblot was washed thrice with TBST for 30 min (10 min for each wash). Immunoblot was incubated with specific antibody against respective protein for overnight in cold condition. After primary antibody probing, immunoblot was washed thrice with TBST for 30 min (10 min for each wash). Membrane was further incubated in either anti-mouse or anti-rabbit horseradish peroxidase (HRP) conjugated secondary antibody (constituted in blocking buffer) for 1 h at room temperature. Membrane was again washed thrice with TBST for 30 min (10 min for each wash). Protein bands were visualized using ECL (chemiluminescent substrate) and chemidoc (UVITEC) instrument.

### Nuclear and cytoplasmic fractionation

For the preparation of cytoplasmic lysate, ice-cold hypotonic cytoplasmic extract buffer (10 mM HEPES, 10 mM KCl, 0.1 mM EDTA and 0.1 mM EGTA) with protease inhibitors was added in the cell pellet and gently mixed with the pipette in a microfuge tube. The cell suspension was incubated on ice for 30 min to allow them to swell. After incubation, freshly prepared 10% NP- 40 was added and vortexed vigorously for 15 seconds to rupture the plasma membrane. The contents were then centrifuged at 13000 rpm for a minute at 4°C and supernatant containing the cytoplasmic lysate was transferred to another pre-chilled microfuge tube and stored at –70°C. The pellet was then further processed for extraction of nuclear lysate. For this, ice-cold nuclear extract buffer (10 mM HEPES, 0.2 M NaCl, 0.5 mM EDTA and 0.5 mM EGTA) with protease inhibitors was added to the pellet and incubated on ice for 45 min with intermittent vortexing after every 10 min of incubation. Finally, cell suspension was centrifuged for 10 min at 14000 rpm. The supernatant containing nuclear lysate was stored at –70°C for further experiment.

### Reverse transcriptase (RT)-PCR

MDA-MB-231 and AMOT stable MDA-MB-231 cells were seeded and grown for 24 h. RNA was isolated by TRIzol method (Gibco BRL, Grand Island, NY) and 5 µg of total RNA was used to synthesise first-strand cDNA using M-MLV RT kit (Invitrogen) and oligo(dT) primer. This first-strand cDNA was used for qPCR for quantification using ThermoScientific Maxima SYBR green/ROX qPCR master-mix (2X) and AMOT or GAPDH specific RT primers. The primer sequence and product size are as follows: AMOT: (190 bp) {forward} 5’- CGTTTGCTACAAGAGCAGCTT-3’; (reverse) 5’-GTTCTTGTCGAGCAGCATGAG-3’; GAPDH: (192 bp) {forward} 5’-ACCTGCCAAATATGATGAC-3’; {reverse}5’- TCATACCAGGAAATG AGCTT-3’.

### Cycloheximide chase assay

MDA-MB-231 and profilin-stable cells pretreated with ATRA (20 µM) for 24 h (Saurav and Manna, 2022a) were incubated with cycloheximide (50 µg/ml) for an ascending time period. Cells were washed with PBS and lysed using NETN buffer followed by immunoblotting.

### Immunoprecipitation

For immunoprecipitation of AMOT with PFN1, HEK293T cells were co-transfected with HA-AMOT and SFB-PFN1 using PEI as a reagent for 24 h. The cells were harvested and lysed in NETN lysis buffer. SFB-PFN1 was precipitated from the cell extracts by incubating with Streptavidin beads overnight. Non-specific interactions were cleared by washing thrice with lysis buffer. Samples were prepared by adding 6× SDS-PAGE Laemmli loading buffer containing β- mercaptoethanol and boiled for 5 min. Immunoprecipitated samples were immunoblotted using rabbit-raised antibodies against HA or PFN1.

### Transwell migration assay

Cell migration was studied using Transwell chambers with 8-µm pore size PET membrane inserts in 24-well plates. Cells were grown, treated and harvested. They were resuspended in serum-free medium and 2.5 × 10^4^ cells in 200 µL was added to the upper chamber. The lower chamber was filled with 600 µL of medium containing 10% fetal bovine serum as a chemoattractant. After incubation at 37 °C with 5% CO2 for 6 h, the non-migrated cells on the upper surface of the membrane were gently removed using a cotton swab. Cells that migrated to the lower surface of the membrane were fixed with 4% paraformaldehyde for 15 min, stained with 0.1% crystal violet for 15 min, and washed with PBS. Migrated cells from at least five random microscopic fields per well were imaged and counted under an inverted light microscope.

### Wound healing assay

For this experiment, the cells were seeded at a density of 3× 10^5^ cells in 60 mm dish and grown in cell culture medium overnight at 24 h of incubation (37°C, 5% CO_2_, humidified atmosphere). This was followed by treatment with ATRA for 24 h and the cells were washed with PBS. Scratches were made using sterile tips, serum free media was added and incubated. Images were taken at different time points under an inverted microscope.

### Gel shift assay of YAP-TEAD of CTGF and OCT-1

To determine the amounts of YAP-TEAD of CTGF, EMSA was conducted essentially as described previously (Raviprakash and Manna, 2014). Briefly, 8–12 μg of nuclear extract (NE) proteins were incubated with ^32^P end-labelled double-stranded CTGF-TEAD and OCT-1 oligonucleotides for 60 min at 37 °C, and the DNA–protein complexes were separated from free oligonucleotides on 6.6% native polyacrylamide gels. The oligonucleotides used in the experiment are: CTGF-TEAD sense 5’-ATGCTGAGTGTCAAGGGGTCAGGATCAA-3’, CTGF-TEAD anti-sense 5’-TTGATCCTGACCCCTTGACACTCAGCAT-3’, OCT1 sense 5’- TGTCGAATGCAAATCACTAGAA-3’ and OCT1 anti-sense 5’- TTCTAGTGATTTGCATTCGACA-3’.

### Molecular docking

The PDB structure of PFN1 (1PFL) was obtained from RCSB protein data bank (Berman et al., 2000). AMOT structures were available in complexes with different proteins and were only partially crystallized. Hence, AMOT structures were generated using Alpha-fold (Jumper et al., 2021) and I-TASSER (Yang and Zhang, 2015). To identify the surface of PFN1 binding to AMOT, and since this is a protein-protein dock, molecular docking was performed using HADDOCK 2.4 (Dominguez et al., 2003; van Zundert et al., 2016). Three individual runs were performed: unbiased, restraint guided docking of actin-binding domain and restraint guided docking of poly-L-proline-binding domain. In the unbiased or blind dock, no active or passive residues were specified, hence HADDOCK generated ambiguous interaction restraints across the full solvent-accessible surface of both molecules. For restraint guided docking, the active interfaces on AMOT (PPXY-1, 239-244), PFN1’s actin-binding domain (Val118, His119, Gly121) and PFN1’s PLP-binding domain (Ser132, His133, Leu134) were defined. Top scoring clusters were analysed for HADDOCK score, population, interface RMSD and buried surface area. Binding energies were calculated by PRODIGY (Xue et al., 2016).

### Statistical analysis

Most of the experiments were done thrice and results are expressed as mean ± SEM. Western blot, EMSA and wound healing images were quantified using ImageJ. Graphs were made using GraphPad Prism version 8.0.2 (GraphPad Software, Inc., CA, USA). Statistical analysis of the samples was made by two-tailed paired Student’s t-test wherever applicable. The P < 0.05 was considered to be significant.

## Results

To explore the role of PFN1, we performed mass spectrometry of PFN1, and found a number of new possible interacting candidates. Among them, we have chosen Angiomotin for further studies. Profilin expression was found to be very low in a triple-negative breast cancer (TNBC) cell, MDA-MB-231. These cells were used for the overexpression of PFN1 and AMOT. Manipulation of these genes and other treatments have not shown any defect in cell growth and cytolysis.

### PFN1 physically interacts with AMOT-p130 in TNBC cells

To ask whether PFN1 engages the Hippo scaffold AMOT, we co-expressed SFB-tagged PFN1 and HA-tagged AMOT-p130 (detailed gene structure is shown in (Fig. 1A) in HEK293T cells and performed streptavidin pull-down. HA-AMOT was recovered in the PFN1 pull-down fraction, demonstrating that the two proteins associate in cells (Fig. 1B).

**Figure 1:**
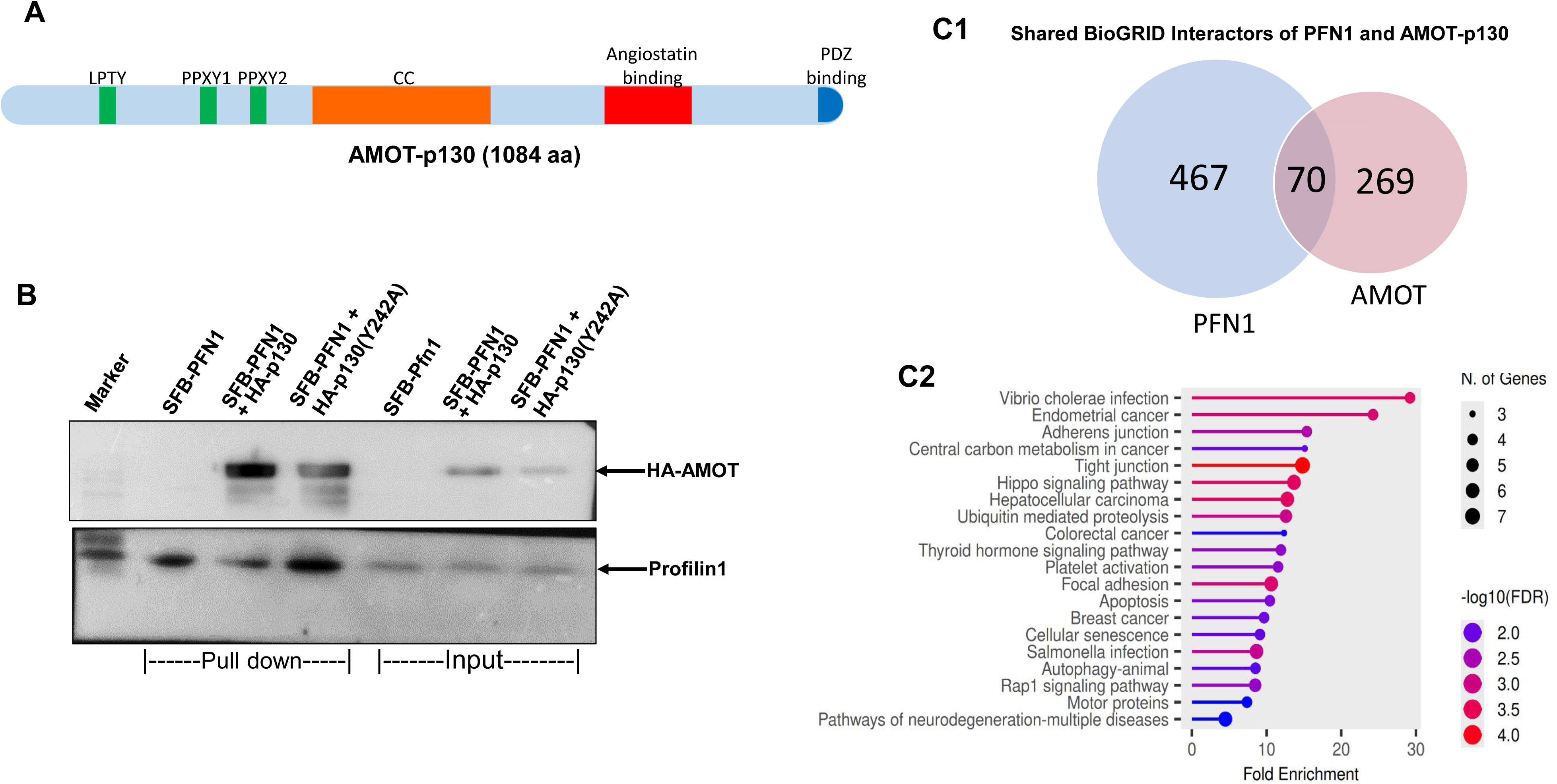
Profilin 1 interacts with AMOT. (A) Domains of AMOT: Coiled coil (CC), Angiostatin binding, PDZ binding. L/PPXY motifs are as indicated. (B) HEK293T cells were transiently co-transfected with HA-tagged Angiomotin and SFB-tagged Profilin1 and the interaction was detected by immunoprecipitation with Streptavidin beads, followed by immunoblotting with anti-HA. (C) Venn diagram of BioGRID-curated physical interactors for PFN1 and AMOT-p130, showing 70 shared interactors between the two interactomes. (D) KEGG pathway enrichment of the 70 shared interactors, ranked by fold enrichment. Dot size denotes the number of genes contributing to each term; color denotes statistical significance (−log10 FDR, scale 2.0–4.0). Pathways of direct relevance to the proposed PFN1–AMOT-p130– YAP1 axis — Hippo signalling, adherens junction, tight junction, and focal adhesion — are among the most significantly enriched terms, along with multiple cancer-associated pathways (endometrial, hepatocellular, colorectal, and breast cancer).

Having established a direct physical interaction between PFN1 and AMOT-p130, we next cross-referenced the curated BioGRID interactomes (Oughtred et al., 2021) of PFN1 and AMOT-p130 and identified 70 shared interactors between the two proteins (Fig.1C1). Among these, the HECT-domain E3 ubiquitin ligase NEDD4 stood out: NEDD4 is an established regulator of AMOT-p130 turnover (C. Wang et al., 2012), and its presence as a shared interactor suggested that the PFN1-AMOT-p130 interaction might intersect with AMOT-p130 protein stability.

Further, KEGG pathway enrichment analysis (Kanehisa and Goto, 2000) of this shared interactor set revealed significant overlap with pathways central to epithelial architecture and growth control, including Hippo signalling, adherens junction, tight junction, and focal adhesion. Several cancer-related pathways were also enriched among the shared interactors, including breast cancer, endometrial cancer, and hepatocellular carcinoma (Fig.1C2).

### PFN1 engages an extended proline-rich region of AMOT independently of the PPxY tyrosines

To define the interaction interface, we performed data-driven protein-protein docking of PFN1 against AMOT using HADDOCK (Dominguez et al., 2003; van Zundert et al., 2016). We performed three independent HADDOCK docking analyses: an unbiased blind dock and two restraint-guided re-docks testing the poly-L-proline (PLP)-binding domain and the actin-binding domain of PFN1 individually (Fig.2A). Despite weaker convergence (top cluster: 4/200 structures), the unbiased dock consistently placed AMOT’s PPxY-1 motif in contact with PFN1’s actin-binding face, anchored by a Glu241(AMOT)–Lys125(PFN1) salt bridge (Fig. 2B). Restraint-guided docking confirmed this preference: the actin-binding domain-restrained run converged far more robustly than the PLP-binding domain-restrained run (57.5% vs. 5.5% of models in the top-scoring clusters), despite near-identical top-cluster HADDOCK scores (−93.8 vs. −94.0). Representative contacts included a Glu241(AMOT)–Arg88/Val118(PFN1) interaction and a Lys245(AMOT)–Glu116(PFN1) salt bridge immediately downstream of the motif (Fig. 2D). Together, these three independent analyses converge on a model in which AMOT interacts with PFN1 predominantly through PFN1’s actin-binding domain rather than the PLP-binding domain.

**Figure 2:**
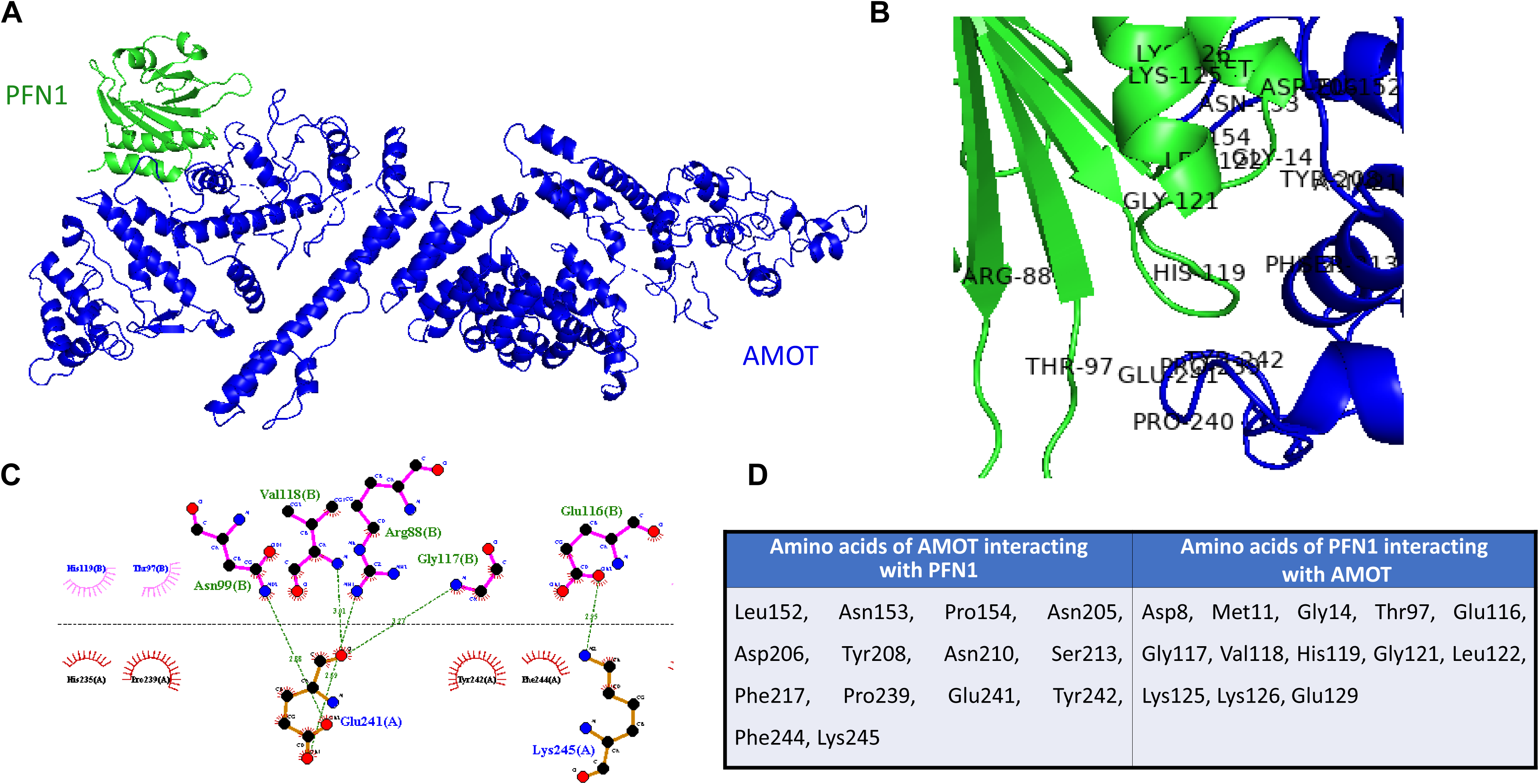
In silico analysis of profilin 1- angiomotin interaction (A) Protein-protein docking was performed to predict the interaction between angiomotin (blue) and profilin1 (green). (B) Amino acids involved in the interaction are indicated. (C) Hydrogen bond interactions between the proteins and the corresponding bond lengths (Å) are shown. (D) The amino acids of angiomotin interacting with profilin1 and the amino acids of profilin1 interacting with angiomotin are listed.

### Generation of AMOT stable cells

As AMOT and PFN1 interacted with each other in transiently AMOT-transfected cells and in silico, the generation of the AMOT stable cells was required to prove this interaction and other biological activities. Detailed panel is shown for transfection and establishment of MDA-MB- 231 breast tumour cells with AMOT stable cells [MDA-MB-231(AMOT^+/+^)] (Fig.5A). The expression of AMOT was increased almost 8-fold as shown by RT-PCR (Fig.5B). The morphology of the cells was unchanged in the AMOT-stable or Profilin-stable MDA-MB-231 cells as compared to wild type cells (Fig.5C). These data suggest the generation of efficient stable cells with AMOT which are going to be used for further experiments.

**Figure 3:**
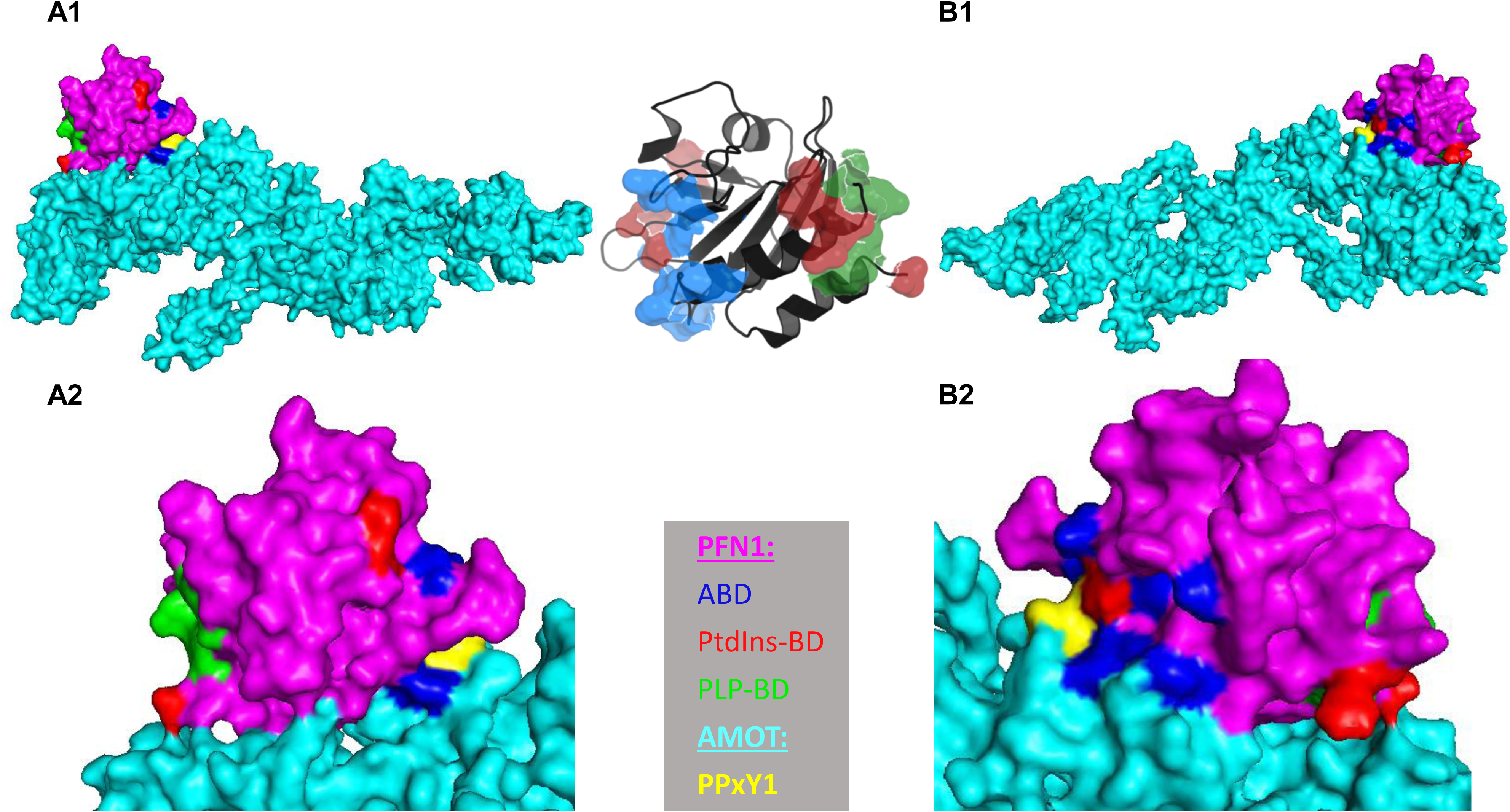
Domains of profilin1 interacting with angiomotin. (A1) Profilin1 (magenta) and angiomotin (cyan) are shown as surfaces. The domains of proflin1 are actin binding (blue), phospho-inositide binding (red) and poly-l-proline binding (green). (A2) zoomed image of (A1) showing actin binding domain at the interface of the interaction. (B1) Horizontally flipped image of the interaction showing the other domains of profilin1. (B2) zoomed image of (B1) showing that the poly-l-proline binding domain (green) is not involved in the interaction.

**Figure 4:**
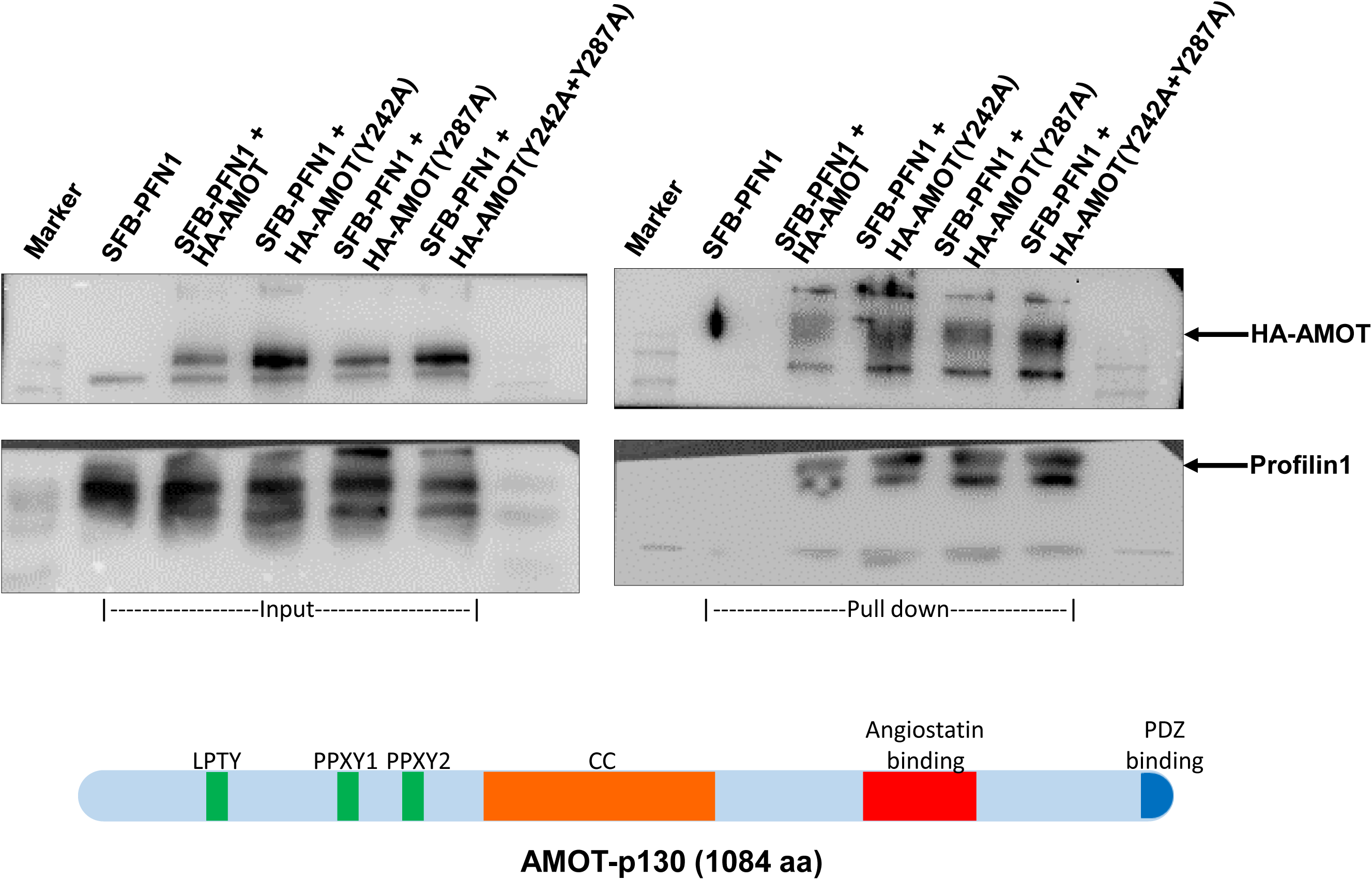
(A) Different mutations of PPXY motifs of angiomotin were co-transfected with SFB- profilin1 and the interaction was determined by incubation with streptavidin beads and was followed by immunoblotting with anti-HA. (B) Different mutations of domains of profilin1 were co-transfected with HA-angiomotin and the interaction was determined by incubation with streptavidin beads and was followed by immunoblotting with anti-HA.

**Figure 5:**
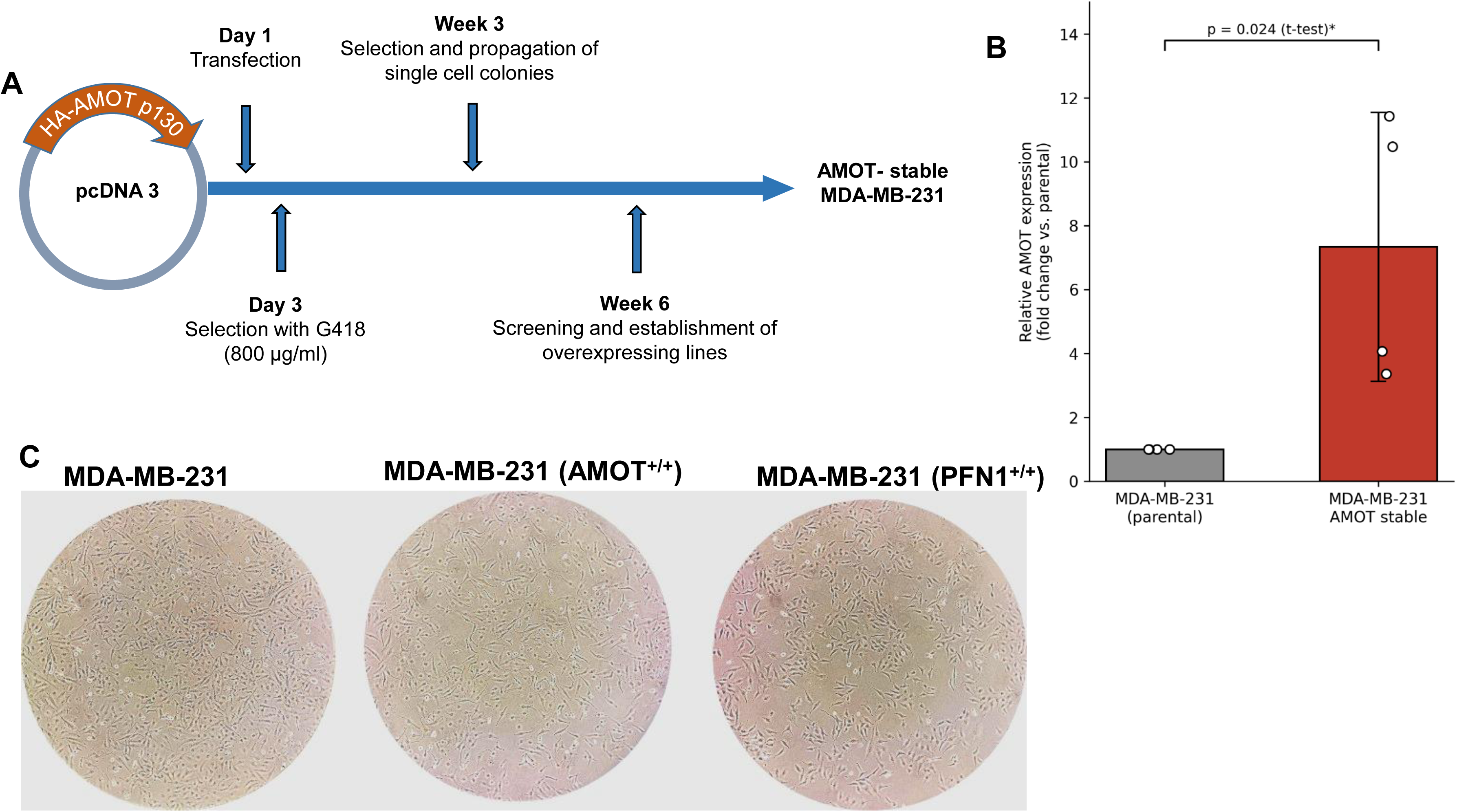
Establishment of AMOT-stable MDA-MB-231 cell lines. (A) Stably overexpressing angiomotin is generated in MDA-MB-231 cells as per described protocol. A work flow of stable cell line generation is described. (B) RT-PCR for angiomotin was performed to confirm the expression levels of angiomotin in the stable cells. GAPDH was used as a control for RTPCR. (C) Phase contrast images were captured to understand the morphology of parental, angiomotin stable and profiling stable MDA-MB-231 cells.

### ATRA stimulation promotes cytoplasmic arrest of YAP in MDA-MB-231(AMOT^+/+^) cells

All-trans-Retinoic acid (ATRA) is a known inducer of PFN1 (Saurav and Manna, 2022a). Amount of PFN1 was increased upon ATRA stimulation in both cells after 24 h of treatment. Sphingosine-1-phosphate (S1P) is a known inducer of YAP (Yu et al., 2012). YAP was induced by S1P in both the cell lines after 12 h of treatment. To evaluate the effect of PFN1 on the translocation status of YAP, MDA-MB-231 and MDA-MB-231(AMOT^+/+^) cells were treated with Sphingosine-1-phosphate (S1P) and ATRA as indicated, and the nuclear and cytoplasmic fractions were analysed by Western blot. Decreased amount of YAP was observed in the nuclear fraction of MDA-MB-231(AMOT^+/+^) cells upon treatment with ATRA (Fig.6A1). The experiments were repeated thrice and amounts of YAP were represented as mean±SEM (Fig.6A2). This result suggests the arrest of YAP in cytoplasm upon AMOT expression.

### Reduced TEAD–DNA binding activity at the CTGF promoter response element

To further understand the effect of PFN1 on arresting YAP, gel shift assay was performed to detect the levels of TEAD, the transcription factor co-activated by YAP. MDA-MB-231 and AMOT stable MDA-MB-231 cells were treated with Sphingosine-1-phosphate (S1P) and ATRA as indicated. In AMOT stable MDA-MB-231 cells treated with ATRA, the level of TEAD was observed to be lowered, confirming the arrest of YAP in this condition (Fig.6B1). The experiments were repeated twice and amounts of TEAD were represented as mean±SD (Fig.6B2).

### Cycloheximide chase assay

To assess the stability of AMOT upon induction of PFN1, MDA-MB-231 and PFN1-stable MDA-MB-231 cells were treated with ATRA (20 μM) for 24 h followed by cycloheximide (50 μg/ml) treatment in a time dependent manner. Since cycloheximide is a known inhibitor of protein synthesis, decreasing levels of AMOT were observed in MB-231 cells, whereas, ATRA pre-treated MDA-MB-231 or profilin-stable cells showed protection of AMOT degradation (Fig.6C).

**Figure 6:**
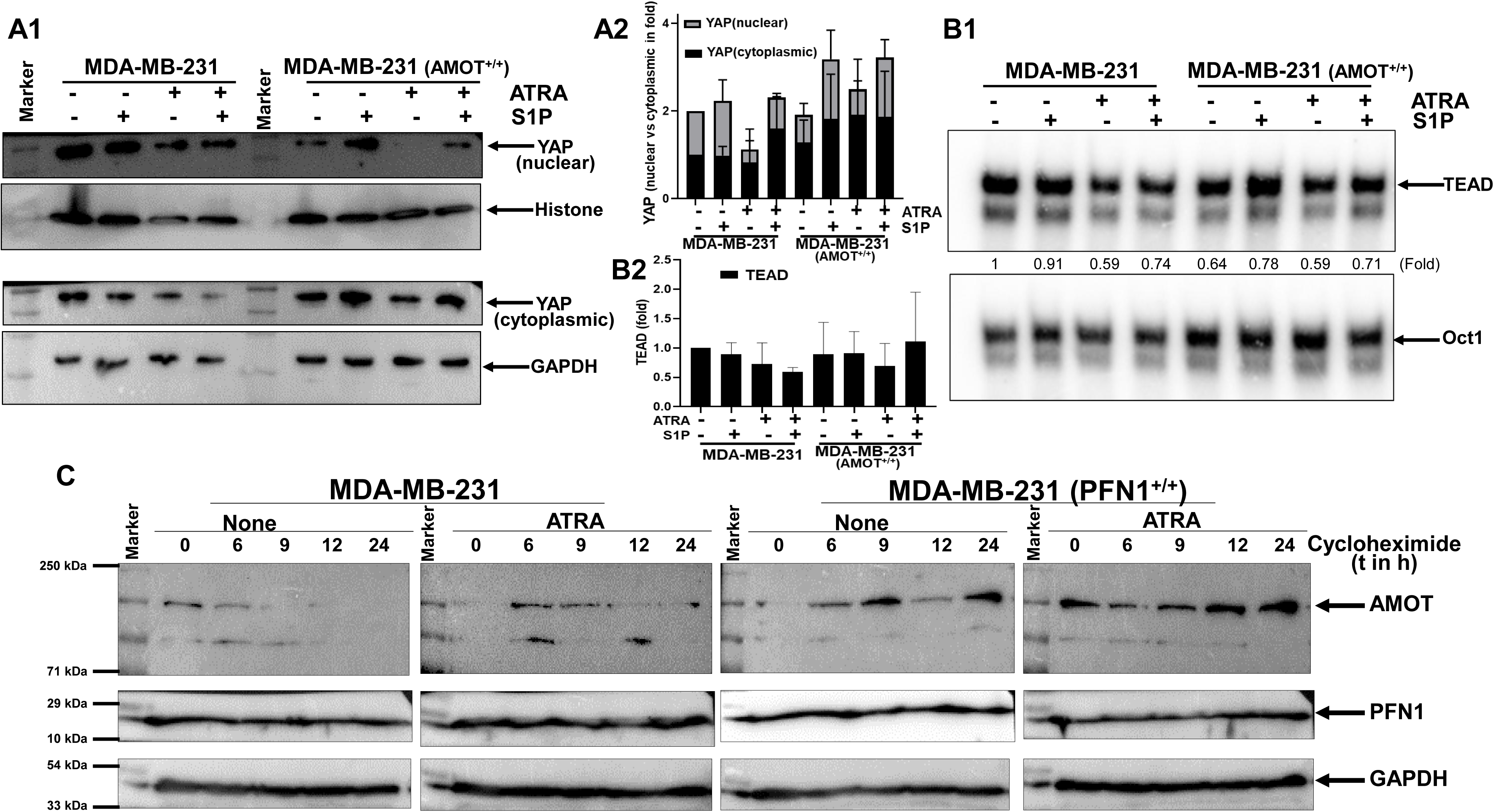
Profilin inhibits YAP nuclear translocation and stabilizes AMOT. (A1) MDA-MB-231 and AMOT stable MDA-MB-231 cells were treated with Sphingosine-1-phosphate (S1P) and ATRA as indicated and nuclear and cytoplasmic fractions were analyzed by Western blot. (A2) The experiments were repeated thrice and amounts of YAP were represented as mean±SEM. (B1) Gel shift assay showing the levels of TEAD when MDA-MB-231 and AMOT stable MDA- MB-231 cells were treated with Sphingosine-1-phosphate (S1P) and All Trans Retinoic acid (ATRA) as indicated. (B2)The experiments were repeated twice and amounts of TEAD were represented as mean±SD. (C) Wild type and profilin-stable MDA-MB-231 cells were treated with ATRA (20 μM) for 24 h followed by cycloheximide (50 μg/ml) treatment in a time dependent manner. The amounts of AMOT, Profilin and GAPDH were determined by Western blot.

### ATRA stimulated MDA-MB-231(AMOT^+/+^) cells show suppression of cell migration

To study the effect of the interaction of PFN1 and AMOT on controlling cell migration, MDA-MB-231, AMOT-stable and PFN1-stable MDA-MB-231 cells were grown and treated with ATRA as indicated and scratches were made with sterile tips. Phase contrast images were captured at different time intervals as indicated under serum depleted conditions (Fig.7A1). The wound closure was observed to be lower in PFN1-stable and ATRA treated AMOT-stable MDA-MB-231 cells as shown in the graph (Fig.7A2). This indicates the effect of AMOT in the presence of Profilin1 in attenuating cell migration.

To further validate the migration studies, MDA-MB-231, AMOT-stable and PFN1-stable MDA-MB-231 cells were grown and treated with ATRA as indicated and were seeded into transwell migration chambers and images were captured after 6 h (Fig.7B1). Cell migration was observed to be lower in PFN1-stable and ATRA treated AMOT-stable MDA-MB-231 cells as shown in the graph (Fig.7B2). This confirms the effect of Profilin1 in attenuating cell migration.

## Discussion

Profilin1 is G-actin sequestering protein, which maintains cell cytoskeleton and cell shape (Witke, 2004). Apart from this, it also interacts with Poly-L-Proline rich proteins as well as with proteins containing poly-inositides (Ding et al., 2012). Out of these two, Poly-L-Proline binding activity is mostly involved in the interactions, affecting cell growth and survival. PFN1 interacts with several phosphatases and controls cell adhesion, motility and invasion. Previous studies reported that PFN1 interacts with PTEN and Akt through its poly-L-proline (PLP) binding domain independent of its actin-binding domain, thereby rendering stability to the said proteins and also recognizing the additional roles of PFN1 besides the traditional actin binding activity (Das et al., 2009). In this study, we have detected the interaction of PFN1 with AMOT which was observed upon co-immunoprecipitation. AMOT-stable and PFN1-stable MDA-MB-231 cells were generated and also ATRA, a known inducer of PFN1 was used (Saurav and Manna, 2022a). Cycloheximide chase assay was performed and it was observed that AMOT is being stabilized by PFN1. Under PFN1 induced conditions of AMOT-stable cells, YAP was observed to be arrested in the cytoplasm by immunoblotting. Reduced TEAD–DNA binding activity at the CTGF promoter response element was confirmed by electrophoretic mobility shift assay (EMSA). Wound healing and Transwell assays were performed to understand the cell migration status which was observed to be attenuated in PFN1 induced AMOT-stable cells and were comparable to that of PFN1-stable cells. These findings establish a novel PFN1–AMOT–YAP axis in which PFN1, by protecting AMOT from proteasomal degradation, enhances cytoplasmic YAP retention and suppresses TEAD-driven transcription, thereby attenuating breast cancer cell migration.

Angiomotin is one of the proteins that interacted with PFN1 as detected by Mass spectrometry data. AMOT is reported as a novel component that inhibits Yes-associated protein (YAP) oncoprotein of the Hippo pathway (Zhao et al., 2011). Hippo pathway is a novel developmental pathway for tissue homeostasis, which is also known as Salvador–Warts-Hippo pathway (Pan, 2010). Several studies suggest the PFN1’s role in tumour suppression (Zou et al., 2007; Das et al., 2009; Bae et al., 2010). However, PFN1 stable cells did not show any significant defect in cell growth or in morphology as shown (Fig.5C). The AMOT-stable TNBC cells also did not show any difference in morphology from parental cells. Whether any pathway is deregulated due to overexpression these proteins in the TNBC cells needs to detected further. PFN1 recognises many of its non-cytoskeletal partners through poly-L-proline-rich sequences read by its PLP-binding domain - the same class of sequence that surrounds the AMOT-p130 PY motifs and governs both YAP binding and NEDD4-mediated turnover (Ding et al., 2009). Despite this convergence, PFN1 has never been linked to the Hippo-YAP1 pathway. Here we show that PFN1 binds AMOT-p130 through its proline-rich region, that this interaction does not depend on the PPxY tyrosines that WW-domain proteins require, that it protects AMOT-p130 from ubiquitin-dependent degradation, and that it restrains YAP1 nuclear accumulation, CTGF transcription and TNBC cell migration. We map PFN1’s binding site on AMOT-p130 and show that it leaves the LPTY motif and both coiled-coil domains free, and we use this to develop a specific model for how PFN1-dependent AMOT-p130 stabilization translates into YAP1 cytoplasmic retention.

Tumour cells’ migration is the hallmark for metastasis. In solid tumour, the cells used to come out of the matrix, pass through the blood and grafted in other organs is regulated by several factors and events. YAP-TEAD mediated expression of several proteins are the important components for these events. Arresting of YAP at cytoplasm by AMOT through its stabilization by interacting profilin is the novel mechanism which is arresting cells’ migration. The wound healing and Transwell migration data (Fig.7) suggest the TNBC cells are affected for cell migration upon AMOT and/or PFN1 overexpression. Several factors are corroborating these events are needs to be detected to understand the deregulation of these molecules upon AMOT and PFN1 overexpression.

**Figure 7:**
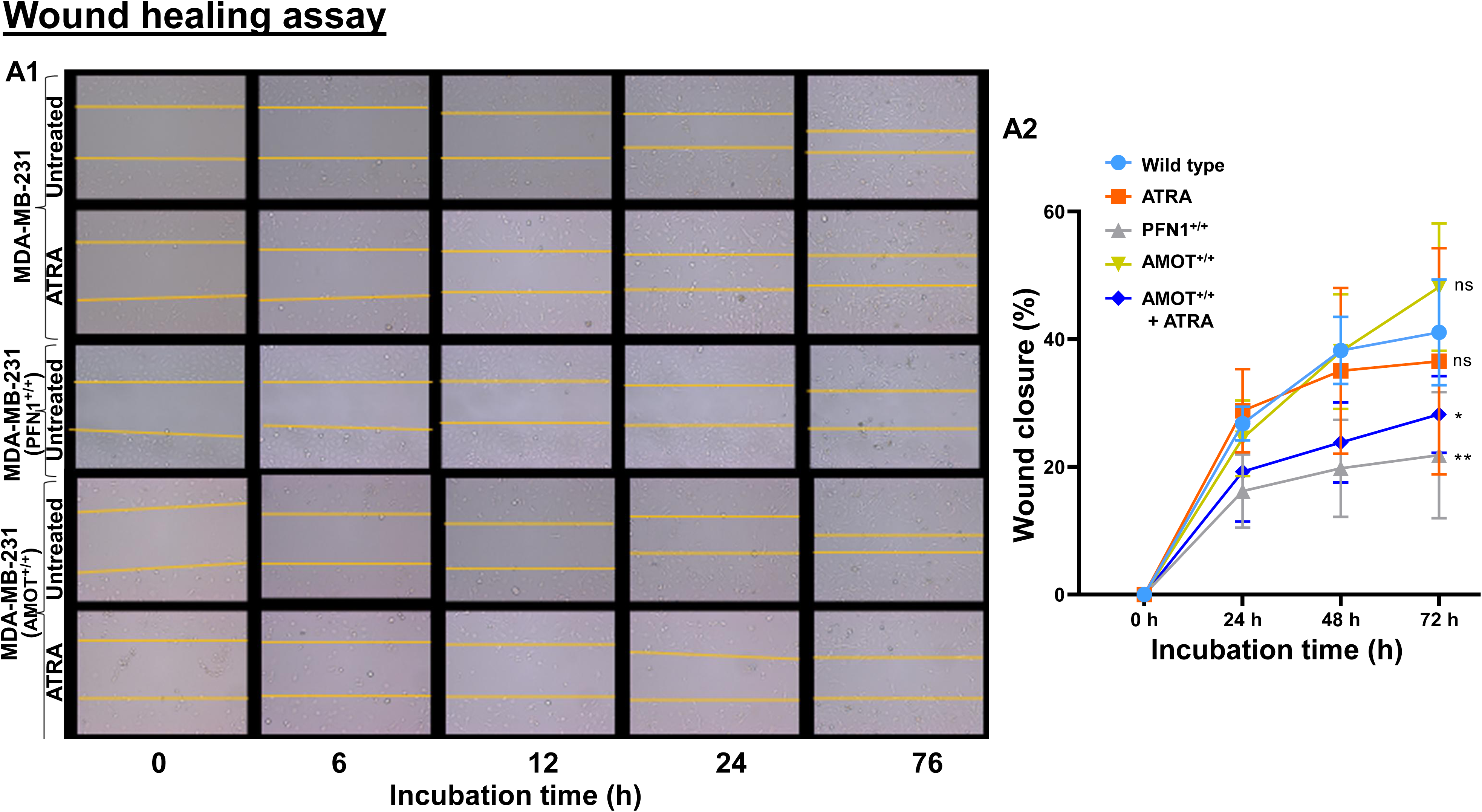
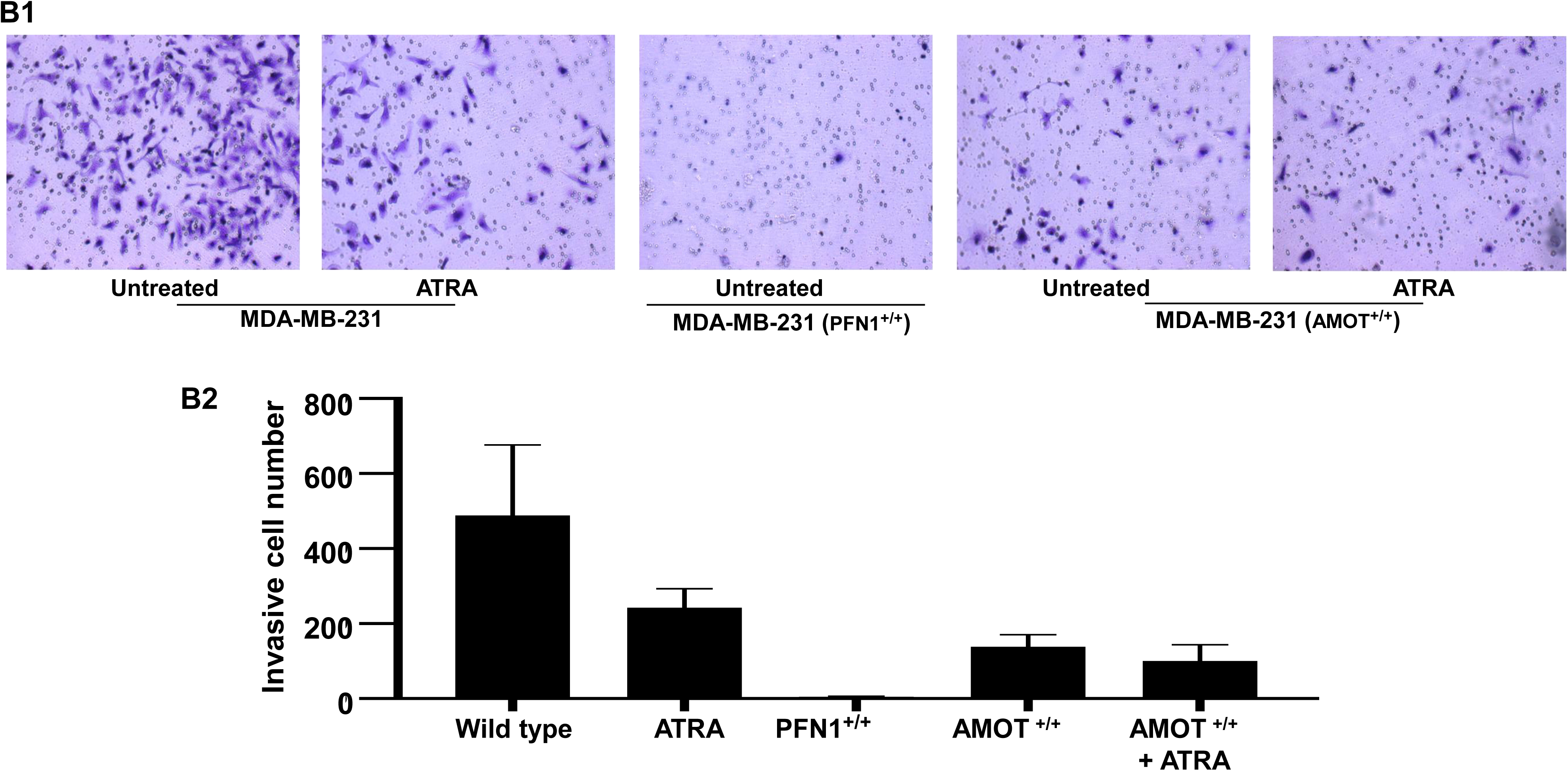
Profilin inhibits tumour cells’ migration. (A1) MDA-MB-231, AMOT-stable and PFN1-stable MDA-MB-231 cells were grown and treated with ATRA as indicated and scratches were made with sterile tips. Phase contrast images were captured at different time intervals as indicated under serum depleted conditions. (A2) Wound closures are indicated in the line diagram in percentage. (B1) MDA-MB-231, AMOT-stable and PFN1-stable MDA-MB-231 cells were grown and treated with ATRA as indicated and were seeded into Transwell Cells’ migration chambers and images were captured after 6 h. (B2) Number of migrated cells were indicated in bar diagram.

Finally, this study indicates an important avenue that Profilin-Angiomotin axis is important to regulate the triple negative breast cancer metastasis. We have shown previously that ATRA is the potent inducer of PFN1 in TNBC and it represses tumour potently with 20 times less concentration of chemotherapeutic agent Vinblastine both in ex vivo and in vivo (Saurav and Manna, 2022a). So, manipulation of profilin by ATRA and increasing the amount of Angiomotin would be a viable strategy to regulate the aggressive triple negative breast tumour metastasis for effective therapeutic.

## Acknowledgement

We thank CDFDs’ Animal House Facility and Sophisticated Equipment Facility (SEF) for its support to finish this study.

## Authors’ contribution

CPV did all the experiments, data curation, investigation, analysis, manuscript writing and SKM conceptualised the project, supervised and wrote the manuscript.

## Abbreviations

AMOT: angiomotin
ATRA: all-trans retinoic acid
CHX: cycloheximide
GAPDH: glyceraldehyde-3-phosphate dehydrogenase
PFN1: profilin 1
TEAD: transcriptional enhanced associate domain
TNBC: triple negative breast cancer
YAP: yes-associated protein.

## References

1. Andrade D, Mehta M, Griffith J, Panneerselvam J, Srivastava A, Kim TD, Janknecht R, Herman T, Ramesh R, Munshi A. YAP1 inhibition radiosensitizes triple negative breast cancer cells by targeting the DNA damage response and cell survival pathways. Oncotarget. 2017;8(58):98495–98508.

2. Bae YH, Ding Z, Das T, Wells A, Gertler F, Roy P. Profilin1 regulates PI(3,4)P2 and lamellipodin accumulation at the leading edge thereby influencing motility of MDA-MB-231 cells. Proc Natl Acad Sci U S A. 2010;107(50):21547–21552.

3. Bergin ART, Loi S. Triple-negative breast cancer: recent treatment advances. F1000Res. 2019;8:F1000 Faculty Rev-1342.

4. Berman HM, Westbrook J, Feng Z, Gilliland G, Bhat TN, Weissig H, Shindyalov IN, Bourne PE. The Protein Data Bank. Nucleic Acids Res. 2000;28(1):235–242.

5. Boopathy GTK, Hong W. Role of Hippo Pathway-YAP/TAZ Signaling in Angiogenesis. Front Cell Dev Biol. 2019;7:49.

6. Bray F, Laversanne M, Sung H, Ferlay J, Siegel RL, Soerjomataram I, Jemal A. Global cancer statistics 2022: GLOBOCAN estimates of incidence and mortality worldwide for 36 cancers in 185 countries. CA Cancer J Clin. 2024;74(3):229–263.

7. Chan SW, Lim CJ, Chong YF, Pobbati AV, Huang C, Hong W. Hippo pathway-independent restriction of TAZ and YAP by angiomotin. J Biol Chem. 2011;286(9):7018–7026.

8. Das T, Bae YH, Wells A, Roy P. Profilin-1 overexpression upregulates PTEN and suppresses AKT activation in breast cancer cells. J Cell Physiol. 2009;218(2):436–443.

9. Ding Z, Bae YH, Roy P. Molecular insights on context-specific role of profilin-1 in cell migration. Cell Adh Migr. 2012;6(5):442–449.

10. Ding Z, Gau D, Deasy B, Wells A, Roy P. Both actin and polyproline interactions of profilin-1 are required for migration, invasion and capillary morphogenesis of vascular endothelial cells. Exp Cell Res. 2009;315(17):2963–2973.

11. Dominguez C, Boelens R, Bonvin AMJJ. HADDOCK: a protein-protein docking approach based on biochemical or biophysical information. J Am Chem Soc. 2003;125(7):1731–1737.

12. Jumper J, Evans R, Pritzel A, Green T, Figurnov M, Ronneberger O, et al. Highly accurate protein structure prediction with AlphaFold. Nature. 2021;596(7873):583–589.

13. Kanehisa M, Goto S. KEGG: Kyoto Encyclopedia of Genes and Genomes. Nucleic Acids Res. 2000;28(1):27–30.

14. Moleirinho S, Guerrant W, Kissil JL. The Angiomotins – From discovery to function. FEBS Lett. 2014;588(16):2693–2703.

15. Oughtred R, Rust J, Chang C, Breitkreutz BJ, Stark C, Willems A, et al. The BioGRID database: A comprehensive biomedical resource of curated protein, genetic, and chemical interactions. Protein Sci. 2021;30(1):187–200.

16. Pan D. The hippo signaling pathway in development and cancer. Dev Cell. 2010;19(4):491–505.

17. Paramasivam M, Sarkeshik A, Yates JR 3rd, Fernandes MJ, McCollum D. Angiomotin family proteins are novel activators of the LATS2 kinase tumor suppressor. Mol Biol Cell. 2011;22(19):3725–3733.

18. Pollard TD, Borisy GG. Cellular motility driven by assembly and disassembly of actin filaments. Cell. 2003;112(4):453–465.

19. Raviprakash N, Manna SK. Short-term exposure to oleandrin enhances responses to IL-8 by increasing cell surface IL-8 receptors. Br J Pharmacol. 2014;171(14):3339–3351.

20. Saurav S, Manna SK. Increased expression of Profilin potentiates chemotherapeutic agent-mediated tumour regression. Br J Cancer. 2022;126(10):1410–1420.

21. Saurav S, Manna SK. Profilin upregulation induces autophagy through stabilization of AMP-activated protein kinase. FEBS Lett. 2022;596(14):1765–1777.

22. Small JV, Stradal T, Vignal E, Rottner K. The lamellipodium: where motility begins. Trends Cell Biol. 2002;12(3):112–120.

23. Stüven T, Hartmann E, Görlich D. Exportin 6: a novel nuclear export receptor that is specific for profilin·actin complexes. EMBO J. 2003;22(21):5928–5940.

24. van Zundert GCP, Rodrigues JPGLM, Trellet M, Schmitz C, Kastritis PL, Karaca E, Melquiond ASJ, van Dijk M, de Vries SJ, Bonvin AMJJ. The HADDOCK2.2 web server: user-friendly integrative modeling of biomolecular complexes. J Mol Biol. 2016;428(4):720–725.

25. Wang C, An J, Zhang P, Xu C, Gao K, Wu D, Wang D, Yu H, Liu JO, Yu L. The Nedd4-like ubiquitin E3 ligases target angiomotin/p130 to ubiquitin-dependent degradation. Biochem J. 2012;444(2):279–289.

26. Witke W. The role of profilin complexes in cell motility and other cellular processes. Trends Cell Biol. 2004;14(8):461–469.

27. Xue LC, Rodrigues JPGLM, Kastritis PL, Bonvin AMJJ, Vangone A. PRODIGY: a web server for predicting the binding affinity of protein-protein complexes. Bioinformatics. 2016;32(23):3676–3678.

28. Yamaguchi H, Condeelis J. Regulation of the actin cytoskeleton in cancer cell migration and invasion. Biochim Biophys Acta. 2007;1773(5):642–652.

29. Yang J, Zhang Y. I-TASSER server: new development for protein structure and function predictions. Nucleic Acids Res. 2015;43(W1):W174–W181.

30. Yi C, Shen Z, Stemmer-Rachamimov A, Dawany N, Troutman S, Showe LC, Liu Q, Shimono A, Sudol M, Holmgren L, Stanger BZ, Kissil JL. The p130 isoform of angiomotin is required for Yap-mediated hepatic epithelial cell proliferation and tumorigenesis. Sci Signal. 2013;6(291):ra77.

31. Yu FX, Zhao B, Panupinthu N, Jewell JL, Lian I, Wang LH, Zhao J, Yuan H, Tumaneng K, Li H, Fu XD, Mills GB, Guan KL. Regulation of the Hippo-YAP pathway by G-protein-coupled receptor signaling. Cell. 2012;150(4):780–791.

32. Yu FX, Zhao B, Guan KL. Hippo Pathway in Organ Size Control, Tissue Homeostasis, and Cancer. Cell. 2015;163(4):811–828.

33. Zaidi AH, Manna SK. Profilin–PTEN interaction suppresses NF-κB activation via inhibition of IKK phosphorylation. Biochem J. 2016;473(6):859–872.

34. Zhao B, Li L, Lu Q, Wang LH, Liu CY, Lei Q, Guan KL. Angiomotin is a novel Hippo pathway component that inhibits YAP oncoprotein. Genes Dev. 2011;25(1):51–63.

35. Zou L, Jaramillo M, Whaley D, Wells A, Panchapakesa V, Das T, Roy P. Profilin-1 is a negative regulator of mammary carcinoma aggressiveness. Br J Cancer. 2007;97(10):1361–1371.

